# Motor activity of nonmuscle myosin 2A is a key component of bipolar filament turnover in cells

**DOI:** 10.64898/2026.03.10.710872

**Authors:** Anil Chougule, Tatyana M. Svitkina

## Abstract

Cell contractility plays numerous essential roles in a healthy organism, while its malfunctioning can lead to disease. The ubiquitous actin-dependent motors of the nonmuscle myosin 2 (NM2) family, which includes NM2A and NM2B, are chiefly responsible for cell contractility because of their ability to polymerize into bipolar filaments. Polymerization/depolymerization of NM2 filaments allows cells to quickly reorganize their contractile system according to constantly changing shapes and positions of nonmuscle cells. Bipolar filament depolymerization is known to depend on the C-terminal features of the NM2A heavy chain. Here, we show that the motor activity of NM2A is another key component of NM2A’s depolymerization mechanism, which cooperates with tail-dependent mechanisms to facilitate NM2A turnover in cells and, through copolymerization with NM2B, to reorganize and dynamize NM2B in trans, thus generating a proper intracellular NM2A/NM2B distribution needed for efficient cell migration. Together, we show that NM2A motor activity is a key component of the bipolar filament depolymerization mechanism.

## INTRODUCTION

Cell contractility plays multiple and diverse roles in a healthy organism, while malfunctioning of this system can lead to disease. Contraction in cells is generated by actin-dependent motors of the myosin 2 family, which have unique ability to assemble their unipolar molecules (with motor domains at one end) into bipolar filaments (with motors at both ends). Bipolar filaments can cause contraction by pulling onto oppositely oriented actin filaments. Myosin 2 motor activity resides in the N-terminal globular head of the heavy chain and the adjacent neck region, which binds two different light chains. To form an individual hexameric myosin 2 molecule (referred to as “monomer” later on) the heavy chain dimerizes via parallel coiled-coil association between its long C-terminal tails. Bipolar filament assembly occurs through both parallel and antiparallel association between these coiled-coil domains of myosin 2 monomers (1-4).

In nonmuscle cells, bipolar filaments of nonmuscle myosin 2 (NM2) can be associated with random actin filament networks or form mutually aligned bundles of actin and NM2 filaments, usually called stress fibers. Both of these structures exhibit contractile properties and undergo constant rearrangements during cell motility (5-11). By this reason, NM2 bipolar filaments, in contrast to their striated muscle-specific paralogs, need to constantly depolymerize and polymerize to keep up with constantly changing shapes and positions of migrating cells. Accordingly, NM2 mutations affecting either motor activity or bipolar filament turnover are associated with multiple human disorders (12).

In contrast to relatively well-defined mechanisms controlling NM2 activation and polymerization (1-3), the depolymerization mechanisms of NM2 filaments are not fully defined (1, 3, 13). Disassembly of NM2 bipolar filaments is typically regulated by the C-terminal region of the heavy chain, which contains the nonhelical tailpiece (NHT) at the very C-terminus, phosphorylation sites within the NHT and at the end of the coiled-coil rod, and binding sites for regulatory proteins. These regions are most divergent among the three mammalian NM2 heavy chains. Accordingly, the most common mammalian NM2 paralogs, NM2A (encoded by MYH9 in humans) and NM2B (encoded by MYH10 in humans), exhibit different turnover rates in cells – faster for NM2A and slower for NM2B (14-17). Moreover, through copolymerization into heterotypic bipolar filaments, NM2A can accelerate NM2B dynamics and broaden its intracellular distribution (1, 18, 19). The underlying mechanism involves preferential dissociation of fast-cycling NM2A subunits from heterotypic NM2A/NM2B filaments, so that the remaining NM2B subunits are forced to recycle and redistribute as well (1). The important roles of NM2 depolymerization mechanisms in fine-tuning the overall design of the cellular contractile system necessitates better understanding of the intricate interplay between specific mechanisms of bipolar filament disassembly.

Along these lines, we recently found that the NHT and the heavy chain tail phosphorylation act synergistically in promoting NM2 dynamics (19). Moreover, we found that the NM2 chimeras, in which the motor and tail domains of NM2A and NM2B were swapped, reorganized the cellular contractile system more efficiently when they contained the NM2A motor rather than the NM2B motor, even if they had identical tails (19). Given that NM2A has a faster motor than NM2B (20, 21), these data suggested that NM2 motor activity is another important factor affecting NM2 turnover. Since the roles of NM2 motor activity in cells were typically studied in the context of contractile force generation, and much less in the context of the overall organization and dynamics of the actin-NM2 system in the cell, although such a role has been proposed (22), these data underscore a compelling rationale to better understand the roles of NM2 motor activity as a new regulatory mechanism in the organization and remodeling of the cellular contractile system.

Here, we directly test the role of the NM2A motor in the dynamics of NM2A itself, as well as in the NM2A-dependent remodeling of NM2B in trans. For this purpose, we use NM2A constructs with deleted motor domain, in which the C-terminal disassembly determinants were either kept intact or mutated by deleting the NHT and abrogating phosphorylation by serine-to-alanine substitutions. We show that both motorless NM2A mutants (with either intact or mutated tail) exhibit slower turnover, reduced incorporation into stress fibers and impaired ability to reorganize endogenous NM2B, whereas the motorless mutant with the mutant tail has more severe defects in these functions. Together, our results reveal that the fast motor activity of NM2A, besides playing a major role in contractile force generation (18, 23-25), is also essential for global intracellular organization and dynamics of the contractile system in the cell.

## RESULTS AND DISCUSSION

Disassembly of NM2A filaments is synergistically promoted by the NHT and the C-terminal phosphorylation sites (Ser1943 in the NHT and Ser1915/Ser1916 just upstream of the NHT) in the NM2A heavy chain. The NM2A mutants, in which these disassembly mechanisms were disabled, individually or in combination, exhibited reduced dynamics both in vitro and in cells (19, 26-29). However, overexpression of the mutant NM2A heavy chain, in which both mechanisms were impaired by deletion of the NHT together with S1915A/S1916A substitutions (NM2A-ΔNHT/2SA), only partially inhibited redistribution of endogenous NM2B in COS7 cells, which do not endogenously express NM2A (19). If these mutations were able to completely disable NM2A filament disassembly, the endogenous NM2B distribution in COS7 cells expressing NM2A-ΔNHT/2SA would remain completely intact. Therefore, additional features of NM2A, potentially the motor domain, should control bipolar filament turnover. To directly test this possibility, we engineered motorless NM2A mutants with either intact or mutated heavy chain tails, GFP-NM2A-Δmotor and GFP-NM2A-Δmotor-ΔNHT/2SA, respectively. Wild type GFP-NM2A and GFP-NM2A-ΔNHT/2SA (with an intact motor domain) were used for comparison in some experiments.

### Motorless NM2A constructs exhibit defective copolymerization with NM2B

Deletion of the motor domain from the NM2 heavy chain leads to uncontrolled polymerization of the heavy chain rods into excessively long polymers (30, 31). However, replacement of the motor with a globular protein, HaloTag, led to the formation of morphologically normal bipolar filaments (20). To test whether our GFP-tagged motorless NM2A constructs are also able to form bipolar filaments, and if so whether they are able to copolymerize with NM2B, we expressed these constructs, or wild type NM2A as control, in COS7 cells and analyzed the expressing cells by superresolution structured illumination microscopy (SIM). NM2B in these cells was revealed by immunostaining (Fig. 1).

**Figure 1.**
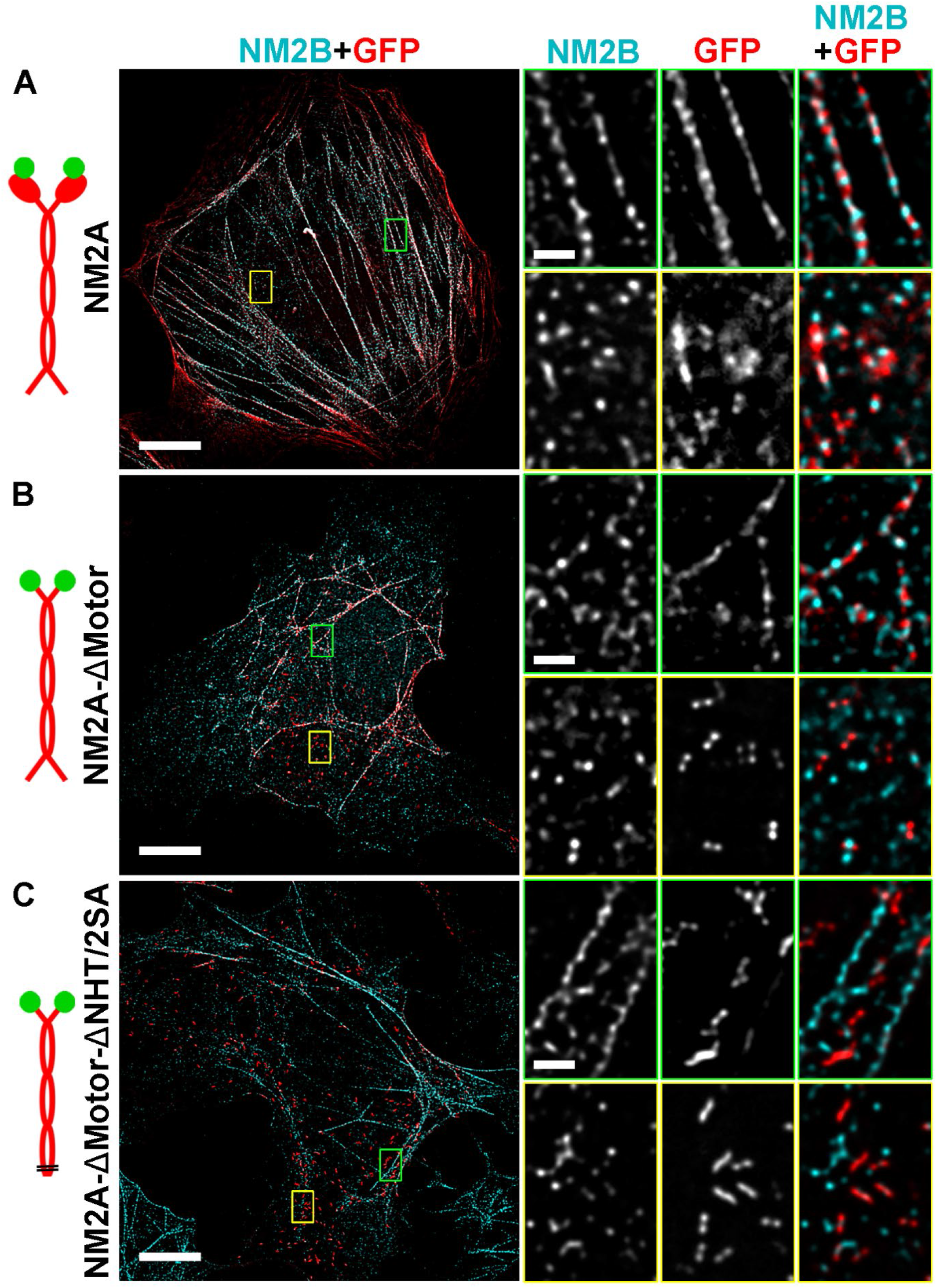
Motorless NM2A forms bipolar filaments that exhibit reduced copolymerization with endogenous NM2B. **(A–C)** SIM images of COS7 cells expressing GFP-NM2A (A), GFP-NM2A-ΔMotor (B), or GFP-NM2A-ΔMotor-ΔNHT/2SA (C) (red in merged images) and immunostained for endogenous NM2B (cyan in merged images). Cartoons (left) illustrate structure of the expressed NM2A constructs including N-terminal GFP tags (green), full-length or truncated NM2A heavy chains (red), and tail mutations (black lines). Boxed regions in main panels are magnified at right to show stress fibers (green boxes) and cytosolic bipolar filaments (yellow boxes). Scale bars: 10 µm (main panels); 1 µm (enlargements).

We found that the expressed GFP-NM2A (shown in red), as expected (32, 33), efficiently copolymerized with endogenous NM2B (cyan) (Fig. 1A). In these cells, virtually all centrally located stress fibers that were positive for NM2B also contained GFP-NM2A. At high magnification, alternating dots of NM2A and NM2B were clearly visible in these stress fibers (Fig. 1A, zoomed region outlined in green). In areas lacking major stress fibers, many individual bipolar filaments could be detected after contrast enhancement. These filaments appeared as red-cyan-red “triple-dot” signals in merged SIM images (Fig. 1A, zoomed region outlined in yellow), which corresponded to GFP-labeled motor domains of NM2A at filament ends (red) and NM2B staining at the filament’s central bare zone (cyan), where the heavy chain tails containing the antibody epitope were located.

The stress fibers at the periphery of the GFP-NM2A-expressing cells were enriched in NM2A. Such pattern is consistent with the typical gradient distribution of NM2A and NM2B in cells that express both paralogs (34, 35). As we showed previously, formation of this NM2A/NM2B gradient is a consequence of copolymerization of NM2A and NM2B combined with their differential dynamics, which results in the displacement of the slow-cycling NM2B from the leading edge by the retrograde actin flow, while the fast-cycling NM2A becomes more broadly distributed and thus more prominent at the cell leading edge (1, 18).

GFP-NM2A-Δmotor expressed in COS7 cells (Fig. 1B) efficiently formed bipolar filaments, which were seen as double red dots in the cytoplasm (Fig. 1B, zoomed region outlined in yellow). These data confirm that GFP, similar to HaloTag (20), can rescue bipolar filament formation in the absence of the motor domain. Importantly, however, only a few of these double-dot red structures contained a cyan dot in the middle, whereas multiple single cyan dots were seen in the area. This pattern suggested that both GFP-NM2A-Δmotor and NM2B could efficiently form homotypic bipolar filaments but copolymerization between them was limited. In stress fibers, copolymerization between GFP-NM2A-Δmotor and NM2B was more evident but still much reduced in comparison with wild type NM2A (Fig. 1A). Copolymerization of GFP-NM2A-Δmotor with NM2B in some stress fibers could be seen at high magnification as alternating red and cyan dots (Fig. 1B, zoomed region outlined in green) and also appreciated based on the whitish shade (overlay of red and cyan) of stress fibers in the cell interior at a lower magnification (Fig. 1B, main panel).

On the other hand, the peripheral stress fibers in these cells were largely cyan in color (Fig. 1B), in contrast to cells expressing GFP-NM2A (Fig. 1A), where peripheral stress fibers had reddish hue. This inverted pattern of NM2B and NM2A-Δmotor distribution, as well as their reduced but not abrogated copolymerization, as compared with wild type NM2A, suggested an important role of the motor in the crosstalk between the two paralogs. These findings are consistent with the described above model of self-organization of NM2 paralogs, wherein more dynamic NM2 versions gain a broader intracellular distribution as compared to less dynamic ones (1, 18). Similarly, a recent preprint study showed that various NM2A mutants and chimeras, which were impaired either in filament disassembly or motor activity, were also shifted away from the cell leading edge acquiring more central cellular distribution comparable to that of wild type NM2B (36).

The defects in copolymerization and intracellular distribution exhibited by GFP-NM2A-Δmotor were dramatically exacerbated in the case of GFP-NM2A-Δmotor-ΔNHT/2SA (Fig. 1C). In cells expressing this mutant, bipolar filaments of GFP-NM2A-Δmotor-ΔNHT/2SA mostly remained in the cytoplasm and were barely, if at all, incorporated into stress fibers (Fig. 1C, zoomed region outlined in yellow). Accordingly, stress fibers contained almost exclusively NM2B (Fig. 1C, zoomed region outlined in green) and appeared mostly cyan at low magnification (Fig. 1C, main panel), indicating that this NM2A mutant had virtually no impact on the stress fiber composition in the expressing cells.

Together, we show here that the motor activity of NM2A facilitates its efficient copolymerization with NM2B, incorporation into stress fibers and broad intracellular distribution, whereas the defects exhibited by the motorless NM2A are exacerbated by the mutations in the heavy chain tails that impair disassembly of NM2A filaments by normal cellular mechanisms. These results reinforce the idea that the motor domain, in addition to generating contractile force, plays a critical role in the contractile system organization.

### Motor and Tail Domains Synergistically Regulate NM2A Turnover Dynamics

Deficient copolymerization of GFP-NM2A-Δmotor with endogenous NM2B (Fig. 1B,C) seems counterintuitive. Since the tail of the GFP-NM2A-Δmotor construct remains intact, the tail-targeting depolymerization mechanisms should be able to release GFP-NM2A-Δmotor subunits from filaments and allow them to interact with available NM2B monomers whose pool is apparently not affected. Therefore, the observed reduced copolymerization of motorless NM2A mutants with NM2B suggests that the motor domain contributes to bipolar filament disassembly. An underlying mechanism could involve active movement of an NM2 monomer along an available actin filament away from the “mother” NM2 filament after it was partially released by the tail-targeting mechanisms. If this is the case, then we can expect that the intrinsic turnover of NM2A in cells is impaired not only by mutations in its tail, but also by deletion of the motor.

We tested this idea using fluorescence recovery after photobleaching (FRAP) of GFP-tagged NM2A constructs expressed in COS7 cells (Fig. 2). For this assay, we used wild type NM2A as control, as well as NM2A mutants with the deleted motor and/or mutated tail. For this analysis, we used stress fibers that did not undergo significant remodeling (contracting, fusing, splitting, breaking, etc.) during the 30-60 min imaging period. After bleaching a small region of the stress fiber, we obtained recovery curves using kymographs generated along the line crossing the bleached region of the stress fiber and fitted these curves to single exponential (Fig 2A-D), as described (19).

**Figure 2.**
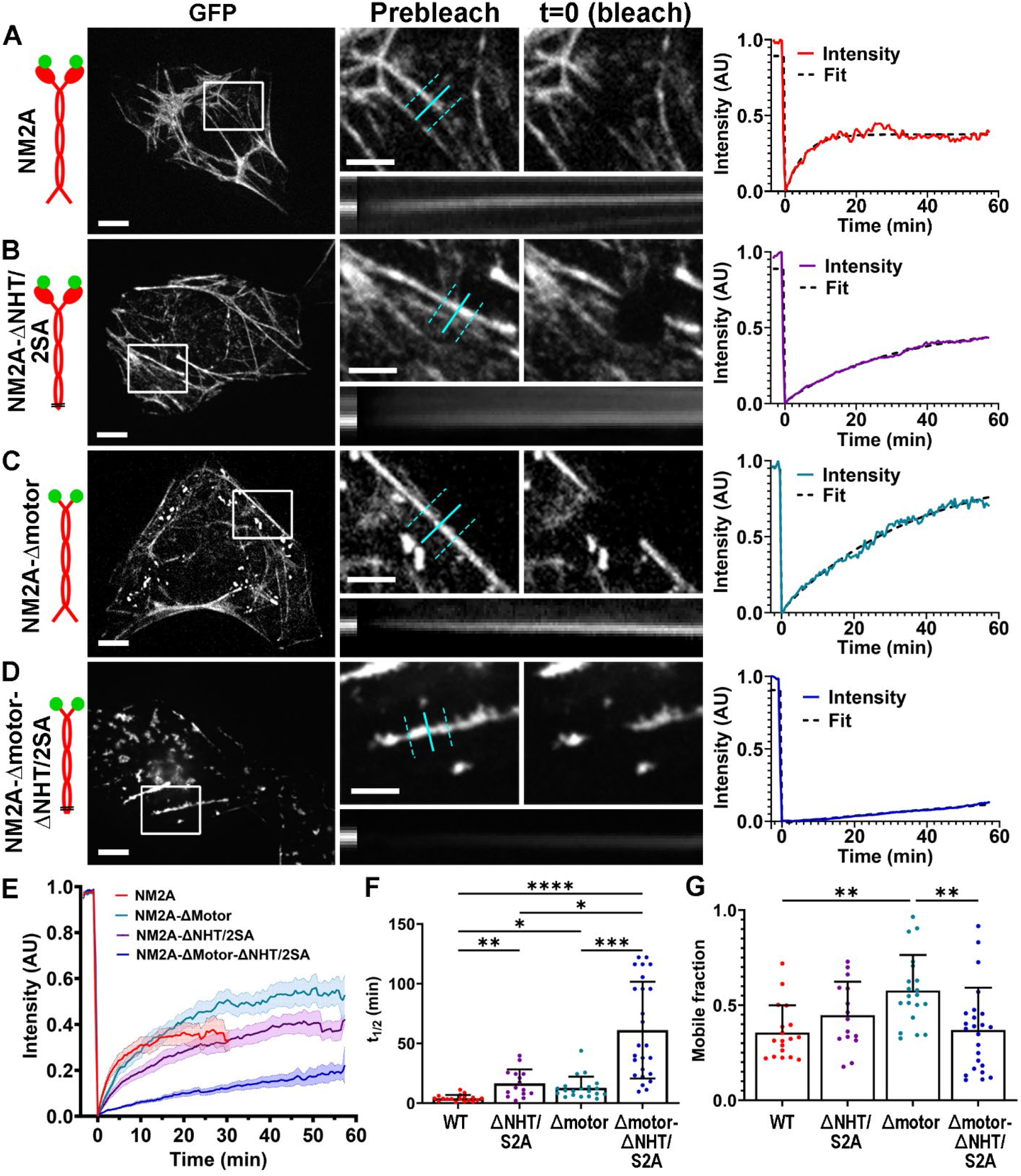
Motor domain accelerates NM2A turnover and synergizes with tail-dependent mechanisms. **(A–D)** Representative FRAP experiments of GFP-NM2A (A), GFP-NM2A-ΔNHT/2SA (B), GFP-NM2A-ΔMotor (C), and NM2A-ΔMotor-ΔNHT/2SA (D) in individual stress fibers of COS-7 cells. Left panels show pre-bleach images with boxed regions indicating areas selected for photobleaching. Upper middle panels show magnified views of boxed regions before bleaching and immediately after bleaching (t = 0). Kymographs (lower middle) were generated along cyan lines, with fluorescence intensity averaged across the region between dashed cyan lines. Right panels show normalized fluorescence recovery curves (colored lines) with single-exponential fits (black dashed lines) for the shown stress fibers. Cartoons (left) illustrate the structure of the expressed NM2A constructs. (E) Mean FRAP recovery curves for GFP-NM2A (red), GFP-NM2A-ΔNHT/2SA (purple), GFP-NM2A-ΔMotor (cyan), and GFP-NM2A-Δmotor-ΔNHT/2SA (blue). Shading indicates SEM. (F, G) Recovery half-times (t_1/2_, F) and mobile fractions (G) for indicated GFP-tagged mutants. Dots represent individual cells; bars indicate means; error bars denote SD. Statistically significant differences are shown as: ****, p < 0.0001, ***, p < 0.001, **, p < 0.01, *, p < 0.1. Scale bars: 20 µm (main panels) and 5 µm (enlargements). The time scale of the kymographs (horizontal direction) is 0-60 min; the kymograph height is 3.3 µm.

By FRAP, GFP-NM2A, as expected, exhibited fast dynamics with an average halftime of recovery (t_1/2_) of 4.2 ± 2.7 min (mean ± SD) (Fig. 2A, E, F). The tail mutations significantly slowed down recovery the GFP-NM2A-ΔNHT/2SA construct resulting in t_1/2_ = 16.7 ± 11.6 min (Fig. 2B, E, F). Remarkably, deletion of the NM2A motor alone in the context of the intact heavy chain tail also significantly reduced the rate of recovery resulting in t_1/2_ = 12.8 ± 9.3 min for GFP-NM2A-Δmotor (Fig. 2C, E, F), which was not statistically different from the value for GFP-NM2A-ΔNHT/2SA. Most impressively, the GFP-NM2A-Δmotor-ΔNHT/2SA construct that lacked both the motor domain and the key regulatory tail determinants exhibited a markedly impaired turnover, with a dramatically prolonged t_1/2_ = 61.2 ± 40.5 min, indicating severe inhibition of filament dynamics, further underscoring the cooperative role of motor activity and tail-dependent regulation in NM2A turnover. The mobile fractions of the NM2A constructs evaluated in these FRAP experiments were mostly similar to each other except for the GFP-NM2A-Δmotor mutant, which exhibited a slightly higher mobile fraction (Fig. 2G). A relatively high mobile fraction of GFP-NM2A-ΔMotor-ΔNHT/2SA shown in Fig. 2G, which seemed inconsistent with the averaged recovery curves (Fig. 2E, blue), likely resulted from extrapolation of exponential fits beyond the actual observation window.

Together, these findings demonstrate that the motor, the NHT, and C-terminal phosphorylation sites are all essential for normal NM2A turnover dynamics and act synergistically to drive NM2A recycling in cells. These data can also explain poor copolymerization of the NM2A-Δmotor-ΔNHT/2SA mutant with endogenous NM2B. Indeed, although newly synthesized NM2A-Δmotor-ΔNHT/2SA monomers should be able to copolymerize with NM2B, the NM2B subunits in such heterotypic filaments can be dissociated by NM2B-dependent mechanisms, whereas the NM2A-Δmotor-ΔNHT/2SA subunits could not. Only if NM2B subunits are present in a large excess, the NM2A-Δmotor-ΔNHT/2SA subunits could be forced to recycle when the NM2B subunits dissociate. This process can explain the observed low level of GFP-NM2A-Δmotor-ΔNHT/2SA turnover. More generally, the mutant subunits would remain bound and gradually recruit additional non-exchangeable NM2A-Δmotor-ΔNHT/2SA monomers until they reach a limit of the filament size (∼30 monomers). The resulting filaments also cannot be incorporated into stress fibers as they lack the actin-binding motor domain. Eventually, homotypic NM2A-Δmotor-ΔNHT/2SA filaments accumulate in the cytoplasm over time, as we observed by SIM (Fig. 1C).

### Motorless NM2A Mutants Fail to Redistribute Endogenous NM2B in COS7 cells

An important role of the NM2A motor domain in the NM2A dynamics raises the possibility that the NM2A motor is also required for efficient redistribution of NM2B in trans, as this process depends on efficient turnover of NM2A filaments (1, 18, 19).

In COS7 cells, which lack endogenous NM2A, the actin-NM2B contractile system consists of a few sharply defined and largely disconnected stress fibers, which do not change their morphology after expression of GFP alone (Fig. 3A). On the other hand, and consistent with our previous findings (1, 18, 19), expression of wild-type NM2A in these cells led to robust redistribution of endogenous NM2B into a more homogeneous pattern, while phalloidin-stained actin filaments reorganized into a system of interconnected and polymorphic stress fibers (Fig. 3B). This crosstalk between NM2A and NM2B also led to a their polarized distribution with NM2A localizing closer to the leading edge than NM2B (Fig. 3B, inset in merged panel), as we reported previously (18).

**Figure 3.**
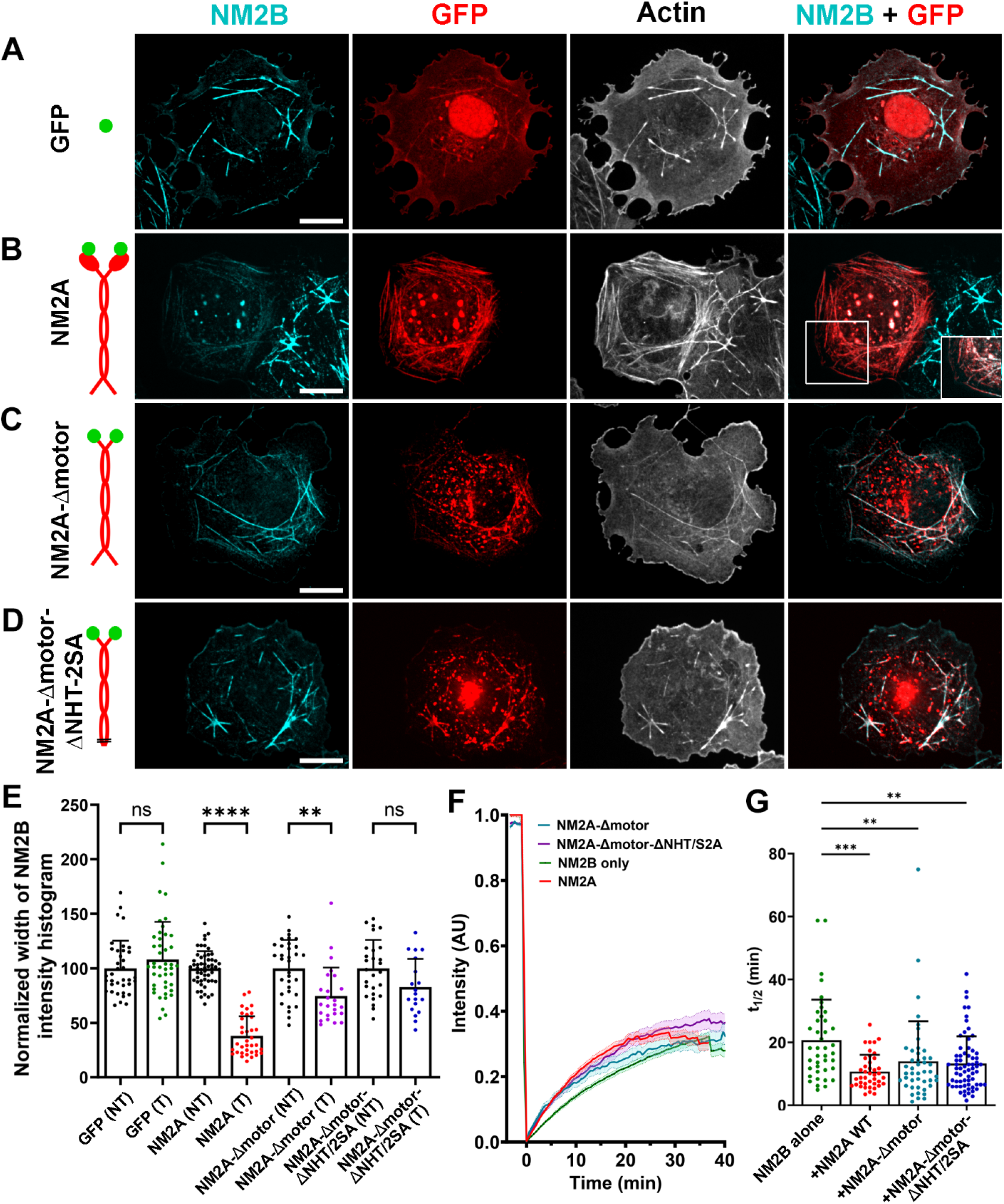
NM2A motor activity is required for redistribution of endogenous NM2B in COS7 cells and synergizes with tail-dependent mechanisms. (A–D) Representative confocal images of COS7 cells expressing GFP alone (A), GFP-NM2A (B), GFP-NM2A-Δmotor (C), or GFP-NM2A-Δmotor-ΔNHT/2SA (D) (GFP is shown in red) and stained with NM2B antibody (cyan) and phalloidin (gray). Merged NM2B and GFP channels are shown in the rightmost panels. Inset in the merged panel B shows the boxed region with enhanced NM2B intensity to show relative enrichment of NM2A at the cell periphery. Scale bars, 20 μm. (E) Distribution of endogenous NM2B in cells transfected with the indicated NM2A constructs (T) quantified based on the width of NM2B immunofluorescence intensity histograms and normalized to the average value of the corresponding non-transfected (NT) cells. Dots represent individual cells; data were collected from at least three independent experiments. Bars indicate mean ± SD. Statistical comparisons were performed using the Kruskal–Wallis test with multiple comparisons; ns, not significant; ****, p < 0.0001; **, p < 0.01. (F) Average FRAP recovery curves of mCherry-NM2B in COS7 cells expressing NM2B alone (green) or coexpressing GFP-NM2A (red), GFP-NM2A-Δmotor (cyan), or GFP-NM2A-Δmotor-ΔNHT/2SA (purple). Shaded regions indicate SEM. (G) Half-times of recovery (t_1/2_) of mCherry-NM2B under the indicated conditions. Dots represent individual cells; bars indicate mean ± SD. ***, p < 0.001; **, p < 0.01.

We quantified the NM2B distribution in cells by measuring the widths of NM2B immunofluorescence intensity histograms, which become narrower for more homogeneous distributions of NM2B, as described previously (18, 19). The average histogram width in expressing cells was normalized against the average value in non-expressing cells from the same coverslip. The results showed that this parameter was not changed by the expression of GFP but was significantly reduced by GFP-NM2A expression (Fig. 3E).

In contrast to wild type NM2A, NM2A-Δmotor induced only a partial redistribution of NM2B, as reflected by a significantly narrower width of NM2B intensity histograms, which however was wider than in cells expressing wild-type NM2A (Fig. 3C, E). Importantly, when the deletion of the motor was combined with the mutations in the heavy chain tail in the GFP-NM2A-Δmotor-ΔNHT/2SA construct, the redistribution of NM2B was largely abolished (Fig. 3D,E).

As a complementary test, we used FRAP to determine whether expression of the NM2A mutants, similar to that of wildtype GFP-NM2A (18, 19), accelerates the dynamics of wild type NM2B. We coexpressed mCherry-NM2B, as a reporter construct, and GFP-NM2A-Δmotor or GFP-NM2A-Δmotor-ΔNHT/2SA, as test constructs (Figure 3F, G). For comparison, we used the previously reported FRAP data for GFP-NM2B and GFP-NM2A, as test constructs (19). The average t_1/2_ of mCherry-NM2B in the presence of GFP-NM2B was 20.7 ± 12.9 min, but it was reduced to t_1/2_ = 10.7 ± 5.4 min in the presence of GFP-NM2A (19). However, both GFP-NM2A-Δmotor and GFP-NM2A-Δmotor-ΔNHT/2SA were less efficient in accelerating the recovery of NM2B, although they still were able to do so resulting in t_1/2_ = 13.9 ± 12.8 min for GFP-NM2A-Δmotor and t_1/2_ = 13.3 ± 8.7 min for GFP-NM2A-Δmotor-ΔNHT/2SA. This limited acceleration of NM2B turnover likely reflects the contribution of a small fraction of copolymers between NM2B and NM2A mutants, which may depolymerize faster due to a smaller number of NM2B subunits within them than in typical NM2B filaments.

These findings demonstrate that the motor domain, in cooperation with the disassembly determinants in the heavy chain tail (NHT and C-terminal phosphorylation sites), not only accelerates NM2A’s own turnover, but also significantly contributes to the turnover of NM2B in trans, thereby changing its intracellular distribution and dynamics.

## CONCLUSIONS

The actin-NM2 contractile system in nonmuscle cells requires efficient reorganization mechanisms in order to keep up with cell migration and shape changes. Regulated depolymerization of NM2 filaments is key part in this process. Our current study reveals a new and rather unexpected component of the NM2A depolymerization mechanism – an important role of the NM2 motor domain, which cooperates with the previously known disassembly mechanisms controlled by the C-terminal region of the NM2A heavy chain (1, 13). On the flip side, our findings also assign a new role to the NM2 motor domain. Although the motor activity of NM2A is typically considered in the context of force generation for the purposes of contraction, tension etc., our data additionally reveal its role in NM2A filament depolymerization.

Using motorless versions of the NM2A heavy chain, we show here that the motor activity of NM2A facilitates its own turnover in cells. In turn, the motor-dependent acceleration of NM2A turnover promotes efficient copolymerization of NM2A with NM2B and, consequently, reorganization and dynamization of NM2B in trans. Through this NM2A/NM2B crosstalk, the cell is able to establish a polarized distribution of NM2A and NM2B, where the fast-cycling NM2A quickly arrives at the new cell front, while slow-cycling NM2B flows backward, where it maintains tension and restricts protrusion (1, 18). In contrast to this normal distribution, we observed an opposite NM2A/NM2B gradient when we used the motorless NM2A mutant (NM2A-Δmotor), suggesting that the fast turnover of NM2A filaments, in part mediated by the NM2A motor activity, also plays an important role in cell-wide organization of the cellular contractile system and distribution of NM2 paralogs, which in turn is important for efficient cell migration (18, 19).

In a broader context, our data suggest that multiple disease-related mutations of NM2A that affect the motor domain may compromise NM2A functions not only by disrupting the proper kinetics of force generation, but also by affecting the turnover of NM2A itself, as well as the dynamics of NM2B in cells, where both paralogs are expressed, leading to problems with cell migration and morphogenesis.

## MATERIALS AND METHODS

### Cells and transfection

Mycoplasma-free COS7 cells (green monkey kidney) were cultured in high-glucose GlutaMAX-DMEM (Gibco) supplemented with 10% fetal bovine serum (Gibco) and 1% penicillin/streptomycin at 37 °C in a humidified incubator with 5% CO_2_. For transfection, cells were seeded onto 35-mm culture dishes and allowed to adhere overnight before transfection using Lipofectamine 3000 (Invitrogen) following the manufacturer’s instructions.

### Plasmids

Plasmids CMV-GFP-NMHC II-A (#11347), CMV-GFP-NMHC II-B (#11348), mCherry-MyosinIIB-C-18 (#55106), and mEGFP-C1 (#54759) were obtained from Addgene. The NM2A-ΔNHT/2SA mutant, which lacks the NHT (amino acids 1928–1961) and carries S1915A and S1916A substitutions, was generated from pCMV-eGFP-NMHC-IIAΔtailpiece (#35689), as described (19). GFP-NM2A-Δmotor and GFP-NM2A-Δmotor-ΔNHT/2SA were generated by replacing the motor domain (amino acids 1-796) with GFP in NM2A WT (#11347) and NM2A-ΔNHT/2SA, respectively. Mutagenesis was performed using the Q5 Site-Directed Mutagenesis Kit (New England Biolabs). Motor-domain deletions were generated using the following primers: Fw 5′-aaagcatttgccaagcggca-3′; Rv 5′-cttgtacagctcgtccatgccgag-3′.

### Immunofluorescence staining

Immunofluorescence staining of NM2B was performed as described previously (19). Briefly, cells were fixed with 4% formaldehyde in PBS, permeabilized using 0.03% Triton X-100 in PBS, and stained with primary rabbit polyclonal antibody (#3404; Cell Signaling) and secondary Alexa Fluor-568 anti-rabbit IgG antibody (#A-11011, Invitrogen). Actin filaments were labeled with Alexa Fluor-647 phalloidin (#8940; Cell Signaling) or iFluor 405-phalloidin (#176752, Abcam).

### Structured Illumination Microscopy (SIM)

Structured illumination microscopy (SIM) was performed using CrestOptics X-Light DeepSIM standalone module mounted on a Nikon Eclipse Ti2E inverted microscope (Nikon Instruments, Japan) equipped with sCMOS camera (Kinetix), CSU-X1 spinning disk (Yokogawa), and Plan Apochromat λD 100×NA 1.45 oil-immersion objective, providing a lateral pixel size of 100 nm (XY). Imaging was carried out using a 488-nm excitation laser with a 510/40 emission filter for GFP and a 545-nm excitation laser with a 595/50 emission filter for Alexa Fluor-568. After acquisition of Z-stack images, image reconstruction was performed using Nikon Imaging Software (NIS-Elements).

### Confocal microscopy

Z-stack images were acquired using Eclipse Ti-U inverted microscope (Nikon Instruments, Japan) equipped with LUNV 7-line laser launch (Nikon), CSU-X1 spinning disk (Yokogawa), QuantEM:512SC CCD camera (Photometrics), and Plan Apo 63×1.4 NA oil-immersion objective and 2× optical zoom and operated by Nikon Imaging Software (NIS-Elements). Excitation was performed using 405-nm, 488-nm, 561-nm, and 640-nm laser lines along with a quad-bandpass filter for simultaneous detection of DAPI, GFP, Alexa Fluor-568 (anti-NM2B), and Alexa Fluor-647 phalloidin fluorescence, respectively.

### Fluorescence recovery after photobleaching (FRAP)

For FRAP experiments, COS7 cells were transfected with either GFP-tagged NM2A constructs only (for figure 2) or cotransfected with GFP-NM2A constructs and mCherry-NM2B (for figure 3). Twenty-four hours post-transfection, cells were replated onto glass-bottom 35-mm Petri dishes and allowed to spread for 6 hours. Prior to imaging, cells were transferred to CO_2_-independent L15 medium for 1 hour at 37°C. During live cell imaging cells were maintained at 37°C in stage-top incubation chamber (Okolab USA Inc.). After capturing single confocal images of GFP and mCherry, if needed, and time-lapse acquisition of 10 pre-bleach frames of the construct of interest (GFP or mCherry), rectangular regions (∼ 16×13 µm) containing stress fibers were bleached for 5 seconds using a 405-nm laser at 100% power. Subsequent imaging of GFP or mCherry was conducted at a single confocal plane using 60×1.4 NA lens at 28.7-second intervals per frame for 1 hour.

### Image analyses

FRAP data analysis was performed using ImageJ software as described previously (19). Briefly, a kymograph generated along a wide (2.2-4.4 μm) line crossing the bleached region of the stress fiber was using to obtain the fluorescence intensity profile over time, which after normalization to the 0 - 1 range was fitted to a single exponential to obtain he half-time recovery (t_1/2_) and the mobile fraction.

Quantification of endogenous NM2B distribution was performed as described previously (18, 19). In brief, the width of NM2B immunofluorescence intensity histograms was measured in 16-bit images using ImageJ. The data from transfected cells were normalized to nontransfected controls from the same samples. Only cells with fluorescence intensity above a set threshold were analyzed to ensure sufficient expression and phenotypic effect.

## Statistical analysis

GraphPad Prism 10.6.1 was used to perform statistical analyses using Kruskal–Wallis test and to generate graphs. Minimum three independent experiments of each type were performed.

## ACKNOWLEDGEMENTS

We thank Justin Bi and Aditya Saha for help with image quantification, and Changsong Yang and Xingyuan Fang for useful discussions. This work is supported by NIH grant R35 GM 140832 to TS.

